# Spatial context allows the evolution the control of nitrification by plants

**DOI:** 10.1101/2024.11.19.624244

**Authors:** Alice Nadia Ardichvili, Sébastien Barot, Jean-Christophe Lata, Nicolas Loeuille

## Abstract

Some plant species inhibit or stimulate soil nitrification, the transformation of ammonium into nitrate by microorganisms. The control of nitrification may in turn alter ecosystem productivity and functioning. Given the potential positive impacts of nitrification control on plant fitness, we aim to determine the conditions under which nitrification control is likely to have been selected, and the consequences of that selection on ecosystem functioning. We investigate both the role of the abiotic context (nutrient availability and diffusion) and the role of other plant traits (mortality and dispersal). A first mean-field model shows that when nitrogen pools are shared among individuals within the plant population, the control of nitrification is counter-selected. A tragedy of the commons occurs because the costs of controlling nitrification (ie. of producing root exudates) only affect the controlling individuals while benefits are shared among all individuals. We then assume that the effects of the control of nitrification are spatially restricted to the rhizosphere, and we build a spatially explicit, individual-based model in which mutation of control of nitrification is possible. Plant capacity to control nitrification evolves when the plant environment is sufficiently private and generation time sufficiently long, leading to higher fitness benefits of the construction process. In such cases, plants evolve to inhibit nitrification when losses of nitrate are greater than losses of ammonium and evolve to stimulate nitrification when losses of ammonium are greater than losses of nitrate. Finally, biomass production tends to be maximal at the selected strategy when the diffusion of ammonium and nitrate is low. Our results help predict which strategies should be selected and likely to be found in different plants in different parts of the world.

## Introduction

Plants are sessile organisms whose growth is constrained by locally available limiting nutrients. The ability of plants to form mutualistic associations with soil microorganisms, such as mycorrhizal associations, the symbiotic fixation of nitrogen (N), and the modification of mineralization rates are examples of such adaptations (Knops et al., 2002). Plant alteration of the N cycle leads to niche construction, as plants modify their environments in a way that alters their own fitness, the fitness of their offspring due to legacy effects, and/or of individuals of other plant species (Odling-Smee et al., 1996).

Several eco-evolutionary models have tackled the evolution of plant modulation of the N cycle and some of its puzzles. Menge and Levin (2017) showed that symbiotic fixation of N could be maintained even in an N-rich system due to spatial heterogeneity. Boudsocq et al. (2011) showed that in some cases plants could evolve a high uptake rate, resulting in over-consumption of the limiting nutrient and leading to the extinction of the population. In fact, a low uptake rate benefits other individuals in the population (leaving more resources available to them) while having an individual cost (reducing individual growth). The evolution of low uptake rates is subjected to the tragedy of the commons (Hardin, 1968) and only evolves when spatial structuring is sufficient (Barot et al., 2016). Spatial structure is also key for the evolution of plant ability to increase mineralization rates (Barot et al., 2014). Increasing mineralization rates or low nutrient uptake rates can be assimilated to altruistic traits. Nitrification, the transformation of ammonium into nitrate by bacteria and archaea, is an important process in the N cycle.

Biological Inhibition of Nitrification (BNI) is the capacity of plants to inhibit the transformation of ammonium into nitrate by microorganisms through the exudation and release of compounds targeting nitrifying microbial communities (Subbarao et al., 2009, 2007, 2015). Such a capacity (and in particular the molecular characterization of compounds responsible of BNI) has been investigated mostly in *Poaceae*. However, empirical evidence also suggests that other plant species, notably trees, may inhibit nitrification (Andrianarisoa et al., 2010; Laffite et al., 2020). Because nitrate is usually more mobile and prone to leaching than ammonium and can be turned into N_2_O by denitrification, plant inhibition of nitrification typically leads to more N-conservative ecosystems, explaining the high productivity of BNI grasses-dominated savannas despite strong N limitation (Boudsocq et al., 2009; Lata et al., 1999). Some plants may also stimulate nitrification (Andrianarisoa et al., 2010; He et al., 2022; McLeod et al., 2016). Carbon-rich root exudates can boost nitrifying communities (Zhang et al., 2019), resulting in higher nitrification rates (Henry et al., 2008). From an ecological point of view, different strategies towards nitrification may explain coexistence between grasses (inhibiting) and trees (stimulating) in savannas (Ardichvili et al., 2024b; Konaré et al., 2019). Though differences between plant species and plant genotypes in their capacity to control nitrification have been documented (Coskun et al., 2017; Lata et al., 1999; Przybylska et al., 2024), the evolution of this capacity has never been thoroughly investigated either empirically or using models.

Under some conditions, plants that control nitrification should experience a more efficient N cycling and have a higher biomass (Ardichvili et al., 2024b); this should result in more resources available for reproduction and higher propagation of controlling phenotypes in the population. The fitness benefits of nitrification control are obvious in two invasive species: the inhibiting *Androbogon gayanus* in Australia and the stimulating *Bromus tectorum* in North America (McLeod et al., 2016; Rossiter-Rachor et al., 2017). Another study showed that the control of nitrification by *Arabidopsis thaliana* is variable and in part genetically determined (Przybylska et al., 2024) suggesting the possibility of evolution of this capacity. Several hypotheses exist as to why the control of nitrification should be selected for (Boudsocq et al., 2009; Lata et al., 2022; Przybylska et al., 2024; Rice and Pancholy, 1972). However, to date, a model describing the evolution of the control of nitrification and testing these hypotheses does not exist. This work aims to investigate the conditions under which the selection of nitrification control is expected, its direction (inhibition or stimulation), and its intensity. Our theoretical results may generate hypotheses regarding the distribution of BNI capacity in savanna grasses at a global scale, which is currently being assessed in a research project (Global Assessment of Nitrification Inhibition by tropical Grasses Project, https://anr.fr/Project-ANR-19CE02-0009).

The control of nitrification modifies biomass production by influencing N losses (Ardichvili et al., 2024b, see above). Whether evolution leads to optimal situations in terms of ecosystem functioning is not given (Metz et al., 2008): first, controlling nitrification may have impacts on other traits. Since the production of root exudates may divert photosynthesis products from growth, the strategy that maximizes individual fitness is subjected to a trade-off and may not correspond to the strategy that maximizes biomass at the population scale. Second, the control of nitrification may benefit neighboring individuals because nutrients may be partly available to them. The extent to which non-controling individuals benefit from the control strategy determines the intensity of the tragedy of the commons and whether evolution can optimize biomass production. We therefore hypothesize that with low costs of the control of nitrification and strong privatization of the mineral N resource, the selected strategy may coincide with the strategy that optimizes biomass production. This broad rationale is in line with work on the evolution of altruism and the tragedy of commons (Ferrière and Le Gaillard, 2001), and models directly addressing the evolution of plant traits influencing nutrient cycling (Barot et al., 2016, 2014).

We use a general N cycling model that we first study in a well-mixed environment, and then simulate the evolution of control in a spatially explicit context. We assume that plant growth is only limited by N, such that more N always results in more plant growth. We investigate two classes of hypotheses: (i) those that constrain the positive or negative evolution of control of nitrification, and (ii) those that influence the direction of control. We hypothesize that the control of nitrification should be selected (i-1) when the longevity of plants is sufficient for them to build their niche, (i-2) when plants have a sufficiently private environment, i.e. when nutrient diffusion to neighboring cells is limited and (i-3) when controlling plants are sufficiently clustered, i.e. when plants disperse over short ranges.

Further, we hypothesize that the direction of the control (inhibition or stimulation) depends on leaching and deposition rates, due to the strong impact of the interactions between the control of nitrification and these parameters on ecosystem functioning (Ardichvili et al., 2024b). We test the hypothesis that (ii-1) inhibition of nitrification should often be selected because ammonium is the first product of mineralization. Boudsocq et al. (2009) suggested that inhibition creates a shortcut in the recycling pathway and should result in a more efficient N cycle. If reducing the number of steps in the N cycle optimizes nutrient cycling, we expect that inhibition of nitrification should be selected for regardless of leaching and deposition rates. On the contrary, we could expect that deposition and leaching rates matter and drive the strength and direction of evolution. We test the hypothesis that (ii-2) the selection of inhibition vs. stimulation should depend on the relative rate of nitrate and ammonium losses from ecosystems, as suggested by Rice and Pancholy (1972); (ii-3) high input rates of ammonium (nitrate) should select for stronger inhibition (stimulation). In addition to being more leachable than ammonium, nitrate is also more mobile in the soil and more likely to diffuse horizontally. Ammonium can be felt as a private resource by plants while nitrate could be a public resource. We expect that this asymmetry in ammonium and nitrate diffusion results in the selection of inhibition (ii-4). Finally, we test the hypothesis that the control of nitrification should to some extent match the plant response trait, ie. that plants preferring ammonium should evolve to inhibit nitrification; plants preferring nitrate to stimulate nitrification (ii-5). Regarding the effect of evolution of ecosystem functioning, we expect that (iii) depending on the level of privatization of the local nitrogen resource, the evolved control of nitrification minimizes, or not, mineral nitrogen losses and maximizes biomass production.

We show that a sufficiently (both spatially and temporally) private environment enables the evolution of the control of nitrification. The selected strategy then depends on leaching rates of ammonium and nitrate, not input rates. Preference for ammonium and nitrate and the propensity of ammonium to diffuse more than nitrate also impact the direction of the selected control, in a way that tends to optimize biomass production when the horizontal diffusion of nutrients is low.

## Methods

### Analytical mean-field model

Our analytical model is derived from a non-spatial recycling model describing N dynamics with 4 compartments: plants *P*, detritus *D*, ammonium *N*_*A*_ and nitrate *N*_*N*_ (Ardichvili et al., 2024b; Boudsocq et al., 2012, Eq. 1, Fig. 1a). We assume that plants are only limited by N and that plant biomass is proportional to plant N content (i.e. that the C:N ratio is fixed). We refer to the size of the plant compartment as plant biomass for simplicity.

**Figure 1:**
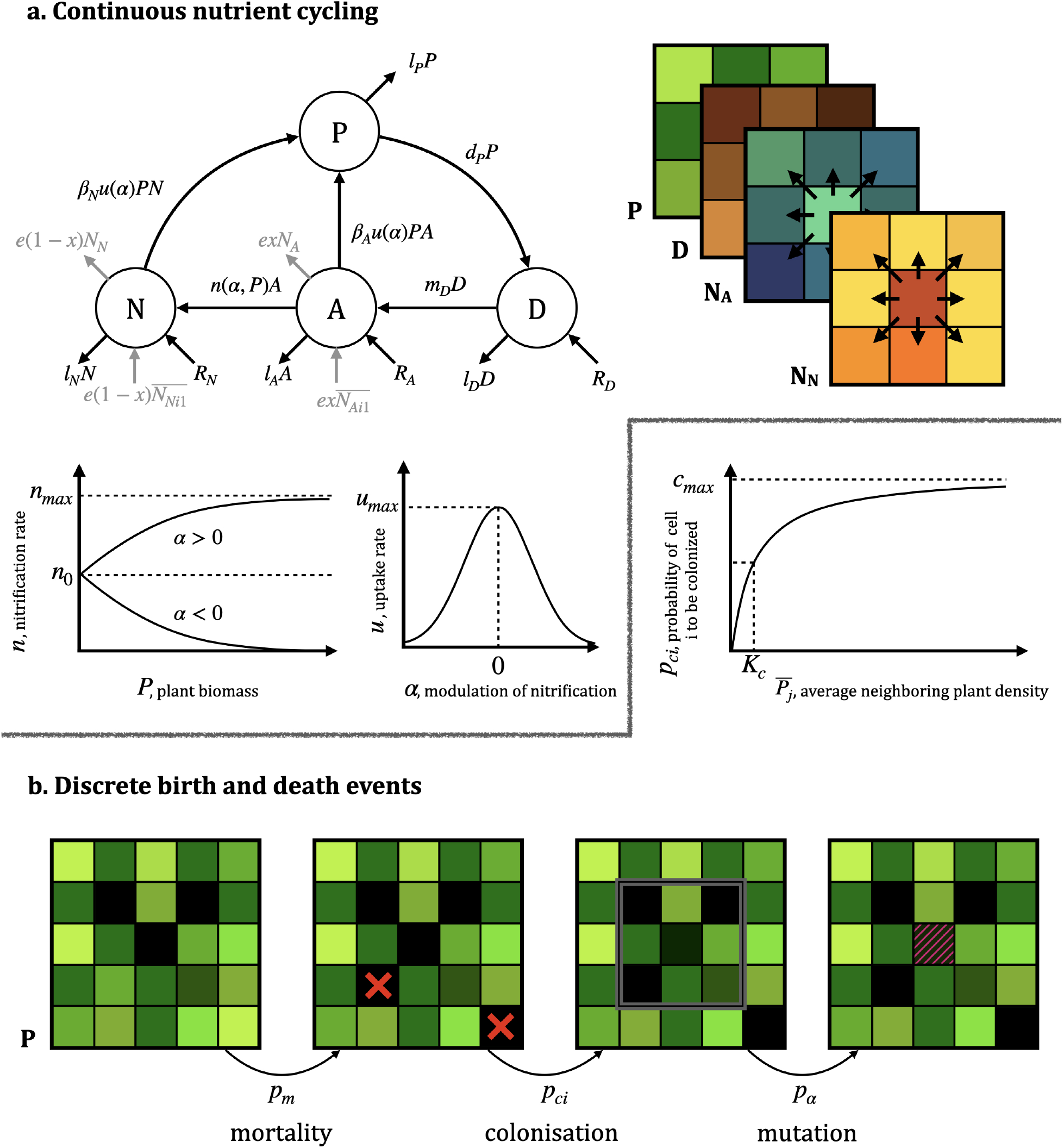
Presentation of the model. Different shades of green, brown, blue, and orange indicate different N content in plants, detritus, ammonium, and nitrate respectively. a. illustrates continuous nutrient cycling dynamics, the black arrows showing the local structure (mean-field model) while the grey arrows show additional fluxes due to spatial dynamics in the spatially explicit simulations b. illustrates the discrete demographic processes. Red crosses indicate where a plant dies (green → black cell). The colonization probability of the middle cell depends on surrounding plant biomass (grey box drawn around the 8 neighboring cells for a dispersion distance of 1 cell). Mutation possibly occurs and changes the strategy of the plant

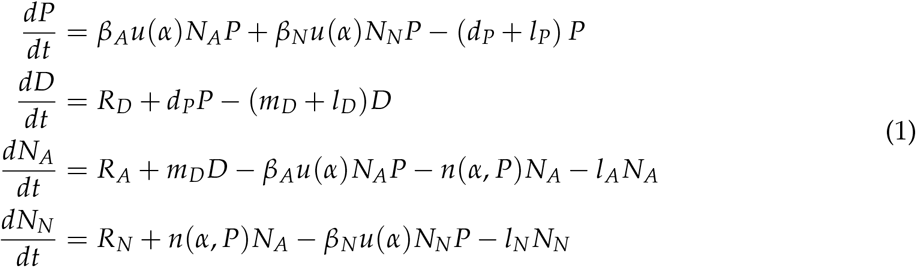

N enters the system via the *N*_*A*_ and *N*_*N*_ pools by atmospheric deposition, or via the *D* pool by detritus import (parameters *R*_*A*_, *R*_*N*_ and *R*_*D*_ respectively). N can also be lost from the *P* compartment at a rate *l*_*P*_ (e.g. due to fire or herbivory), from the *D* compartment at a rate *l*_*D*_ (e.g. because of fire or erosion), from the *N*_*A*_ compartment at a rate *l*_*A*_ (by volatilization), and from the *N*_*N*_ compartment at a rate *l*_*N*_ (by denitrification and leaching). Plants take up N from the ammonium and nitrate compartments at a baseline rate *u* that is modulated by plant preference for ammonium *vs* nitrate (*β*_*A*_/*β*_*N*_, with *β*_*A*_ + *β*_*N*_ = 1). Plant parts die and join the *D* compartment at a rate *d*_*P*_, detritus is mineralized at a rate *m*_*D*_, ammonium is nitrified at a rate *n*, which is modified by the control of nitrification by plants (Fig. 1a):

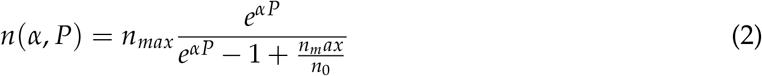

This control depends on per biomass investment into control *α*, the trait of interest in this study and which evolves, and on plant biomass *P*. When there are no plants (*P* = 0) or when plants do not invest in control (*α* = 0), ammonium is nitrified at a constant baseline rate *n*_0_. When *α <* 0, plants inhibit nitrification (BNI), so that *n* decreases with plant biomass, asymptotically reaching 0. When *α >* 0, plants stimulate nitrification and n increases with plant biomass, asymptotically reaching a maximum nitrification rate *n*_*max*_. We assume that the production of root exudates responsible for nitrification control *α* is energetically costly for plants, reducing the uptake rate *u* (Fig. 1a):

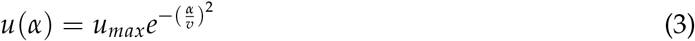

*u* is maximal when plants do not control nitrification (*α* = 0) and decreases as plants inhibit or stimulate nitrification. Parameter *v* determines the strength of the cost of nitrification control - small *v* leading to higher costs. Parameters are provided in Table 1.

**Table 1:** Variables, parameters, units, and default value.

|  | Variables | Unit | Default value |
| --- | --- | --- | --- |
| $t$ | time | yr | - |
| $P$ | patch N content in the plant | $\text{g m}^{-2}$ | - |
| $D$ | patch N content in detritus | $\text{g m}^{-2}$ | - |
| $N_A$ | patch soil ammonium content | $\text{g m}^{-2}$ | - |
| $N_N$ | patch soil nitrate content | $\text{g m}^{-2}$ | - |
| $\alpha$ | strength of control of nitrification | $\text{g}^{-1} \text{m}^2$ | - |
| Cycling parameters |  |  |  |
| $d_P$ | plant recycling rate | $\text{yr}^{-1}$ | 0.6 |
| $l_P$ | plant loss rate | $\text{yr}^{-1}$ | 0.4 |
| $R_D$ | annual inputs of detritus | $\text{g m}^{-2} \text{yr}^{-1}$ | 1.6 |
| $m_D$ | mineralization rate | $\text{yr}^{-1}$ | 0.025 |
| $l_D$ | detritus loss rate | $\text{yr}^{-1}$ | 0.0027 |
| $R_A$ | annual inputs of ammonium | $\text{g m}^{-2} \text{yr}^{-1}$ | 1.5 |
| $l_A$ | ammonium loss rate | $\text{yr}^{-1}$ | 1.25 |
| $R_N$ | annual inputs of nitrate | $\text{g m}^{-2} \text{yr}^{-1}$ | 1.5 |
| $l_N$ | nitrate loss rate | $\text{yr}^{-1}$ | 1.25 |
| $n_0$ | baseline nitrification rate | $\text{yr}^{-1}$ | 2.7 |
| $n_{max}$ | maximum nitrification rate | $\text{yr}^{-1}$ | 4.16 |
| $u_{max}$ | maximum uptake rate | $\text{g}^{-1} \text{m}^2 \text{yr}^{-1}$ | 1.4186 |
| $v$ | cost of nitrification control | $\text{g}^{-1} \text{m}^2$ | 0.5 |
| $\beta_A$ | plant preference for ammonium | - | 0.5 |
| $\beta_N$ | plant preference for nitrate | - | 0.5 |
| $e$ | total inter-cell diffusion | $\text{yr}^{-1}$ | 0.125 |
| $x$ | ammonium propensity to diffuse relative to nitrate | - | 0.5 |
| Demographic parameters |  |  |  |
| $c_{max}$ | maximal colonization probability | - | 1 |
| $d_c$ | plant dispersion distance | - | 1 |
| $K_c$ | average plant density to reach $c_{max}/2$ | - | 3.28 |
| $p_\alpha$ | probability that a mutation occurs | - | 0.01 |
| $\Delta_\alpha$ | amplitude of mutation | $\text{g}^{-1} \text{m}^2$ | 0.005 |
| $p_m$ | probability of plant extinction | - | 0.02 |

In the non-spatial (mean-field) model, we use adaptive dynamics (Dieckmann and Law, 1996; Doebeli and Dieckmann, 2000; Metz, 1992) to analyze the evolution of a plant population characterized by the trait *α* in the absence of spatial structure (Appendix A).

### Spatially explicit model

#### Ammonium and nitrate diffuse to neighboring cells

In a second model, we assume that nitrification control is locally restricted (e.g. within its rhizosphere and immediate surroundings) so that plants controlling nitrification impact their own nutrient cycle. We simulate individual plants on a grid. Each plant grows on a 1m^2^ cell *i*, in which nutrient cycling occurs as in the mean-field model (Eq. 4, 2, 3, Fig. 1). The difference with the mean-field model is that ammonium and nitrate diffuse in neighboring cells at a rate *e*. Neighboring cells are the 8 cells situated in the 8 directions of the focal cell (queen contiguity matrix). We assume that diffusion can be asymmetric between nitrate and ammonium (e.g., to mimic the more usual situation where nitrate is more mobile than ammonium); the strength of this asymmetry is controlled by parameter *x*. We vary *x* continuously between 0 and 1, 0 being the extreme case where only nitrate diffuses, 1 the case where only ammonium diffuses. Let 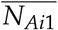 (resp. 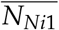) be the average concentration of ammonium (resp. nitrate) in the immediate neighborhood (1 cell away) of focal cell *i*. The nutrient cycling dynamics in a cell *i* are thus described by:

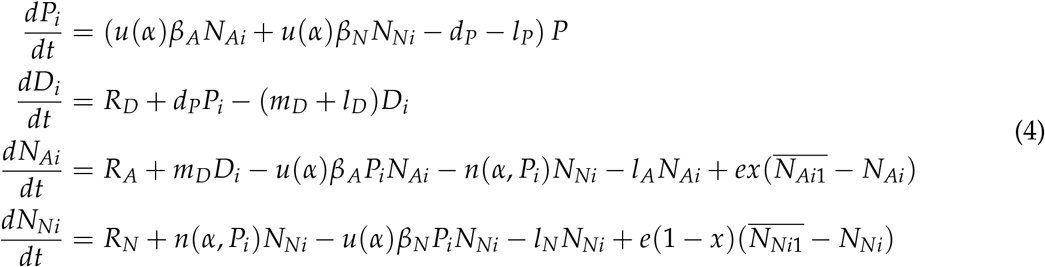

where *n*(*α, P*) is as in Eq. 2, and *u*(*α*) as in Eq. 3.

#### Annual reproduction and mortality

In addition to continuous nutrient dynamics, discrete colonization and extinction events occur every year (Fig.1b). We assume that larger individuals allocate more energy to reproduction and are more fertile. Every cell *i*, if empty, can be colonized with a probability *p*_*ci*_ that depends on the total biomass of surrounding plants (Eq. 5). We write 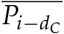 the average biomass of plants situated within a range *d*_*c*_ of the focal cell *i*, where *d*_*c*_ captures the extent to which seeds disperse, i.e. the dispersal strategy of the plant. In Fig. 1b, the plotted *d*_*c*_ range is 1. When colonization is successful, plants inherit a control trait *α* from a parent chosen within the dispersal range, with a probability proportional to the biomass of each potential parent. Mutations occur with probability *p*_*α*_ in which case the seedling trait is slightly higher or lower than the parent trait. Mutations have a fixed amplitude of 0.005.

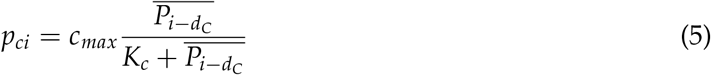

Each year plants die with a probability *p*_*m*_ (due to natural mortality or disturbance). Plant biomass is then transferred to the detritus, so that *D*_*t*+1_ = *D*_*t*_ + *P*_*t*_ and *P*_*t*+1_ = 0. On average, plant individuals die every 1/*p*_*m*_ timestep, defining the plant generation time.

#### Implementation

The simulations are implemented the julia programming language 1.6.0 (Bezanson et al., 2017). The landscape is made of a torus of 50×50 cells. Simulations start with a fully occupied landscape where all compartments are at the local ecological equilibrium (ie. of system (1)). Changing the proportion of occupied cells at the beginning of the simulation did not change the outcome of eco-evolutionary dynamics. Plants initially do not control nitrification (*α*_*ini*_ = 0). For illustration purposes, we also present simulations with a different initial trait to accentuate the direction of evolution. Changing the initial trait did not affect the eco-evolutionary dynamics. The differential equations are integrated using Euler’s method with *h* = 0.05. The simulation stops when *α*, the evolving trait, reaches an equilibrium, i.e. the mean trait and the trait variance over the last 50,000 timesteps do not vary by more than 0.01 and 0.0005, respectively. 10 replicates of each scenario are run.

We first assess whether directional evolution occurs in 4 parameter sets from a previous ecological study (Ardichvili et al., 2024b) corresponding to the Lamto savanna, the Pawnee short-grass prairie, an agricultural landscape, and a toy parameter set with high ammonium volatilization rates and high nitrate deposition. These parameter sets are used to cover a variety of situations, the Lamto ecosystem having large nitrate outflows and being an ecosystem where nitrification control is important (Lata et al., 2004, 1999; Srikanthasamy et al., 2018), the Pawnee ecosystem having much less potential for overall nitrification control. The two other parameter sets are less grounded in empirical observations but allow us to test (i) the role of the asymmetry of fluxes between ammonium and nitrate (toy model); (ii) possible implications from an agricultural point of view. However, many parameters vary between these four parameter sets. We complemented our study by systematically doubling of halving the parameters governing nutrient dynamics *x, l*_*A*_, *l*_*N*_, *R*_*A*_, *R*_*N*_ from a baseline, perfectly symmetric parameter set where *R*_*A*_ = *R*_*N*_, *l*_*A*_ = *l*_*N*_, *x* = 0.5, *β*_*A*_ = *β*_*N*_ = 0.5. Similarly, to test the effect of plant traits, we choose two situations in which inhibition and stimulation evolve (with *β*_*A*_ = 0.2, a situation leading to the evolution of stimulation) and then categorically vary parameters that determine how private the local environment is for the plant (variations of *e, d*_*C*_), and its generation time (variations of *p*_*m*_).

## Results

### No evolution of control in the mean-field model

Assuming no spatial structure, the invasion fitness of a mutant, that is, the *per capita* growth rate of a rare mutant with trait *α*_*m*_ in the stationary conditions set by the resident population of trait value *α* (Metz, 1992), is equal to (see Appendix A for details) :

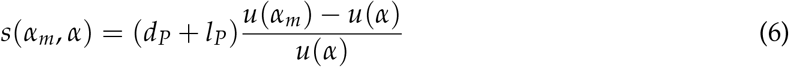

Because mutants replace residents only when the invasion fitness is positive, mutants are selected as long as their basic uptake rate is higher than the basic uptake rate of the resident. Therefore, evolution here leads to the optimization of *u*(*α*), to *α* = 0. Eq. 6 implies that with any preference, for any environmental parameters, the control of nitrification is therefore always counter-selected in a mean-field model.

### Spatial context allows the evolution of nitrification control by plants

Fig. 2 shows that the control of nitrification can evolve when spatial structure is accounted for. Starting with a stimulating plant (at year 0), repeated mutations and subsequent invasions yield the temporal dynamics shown by Fig. 2. The direction of selection and resulting selected control traits depend on ecosystem types. Pale points show all phenotypes present in the grid at a given time. They indicate that a variability of traits coexists around the mean value at all times. Tendencies toward divergent strategies seem to emerge (for example between years 200000 and 250000 in the toy parametrization), but these tentative diversifications are rapidly lost. The mean values of the selected control often follow our predictions. With the parametrizations “Lamto” and “cultivated” (solid orange line and dotted green line in Fig. 2) where ammonium losses are weaker than nitrate losses, ammonium is not only first on the cycle, but also a particularly profitable resource. Inhibition of nitrification is selected. With the toy parametrization (purple dashed line) where there are low nitrate losses and high nitrate deposits, nitrate is a particularly profitable resource and accordingly, the selected strategy is strong stimulation. In the Pawnee prairie (dashed blue line in Fig. 2) where inputs of ammonium and nitrate are balanced, the selected strategy is no control.

**Figure 2:**
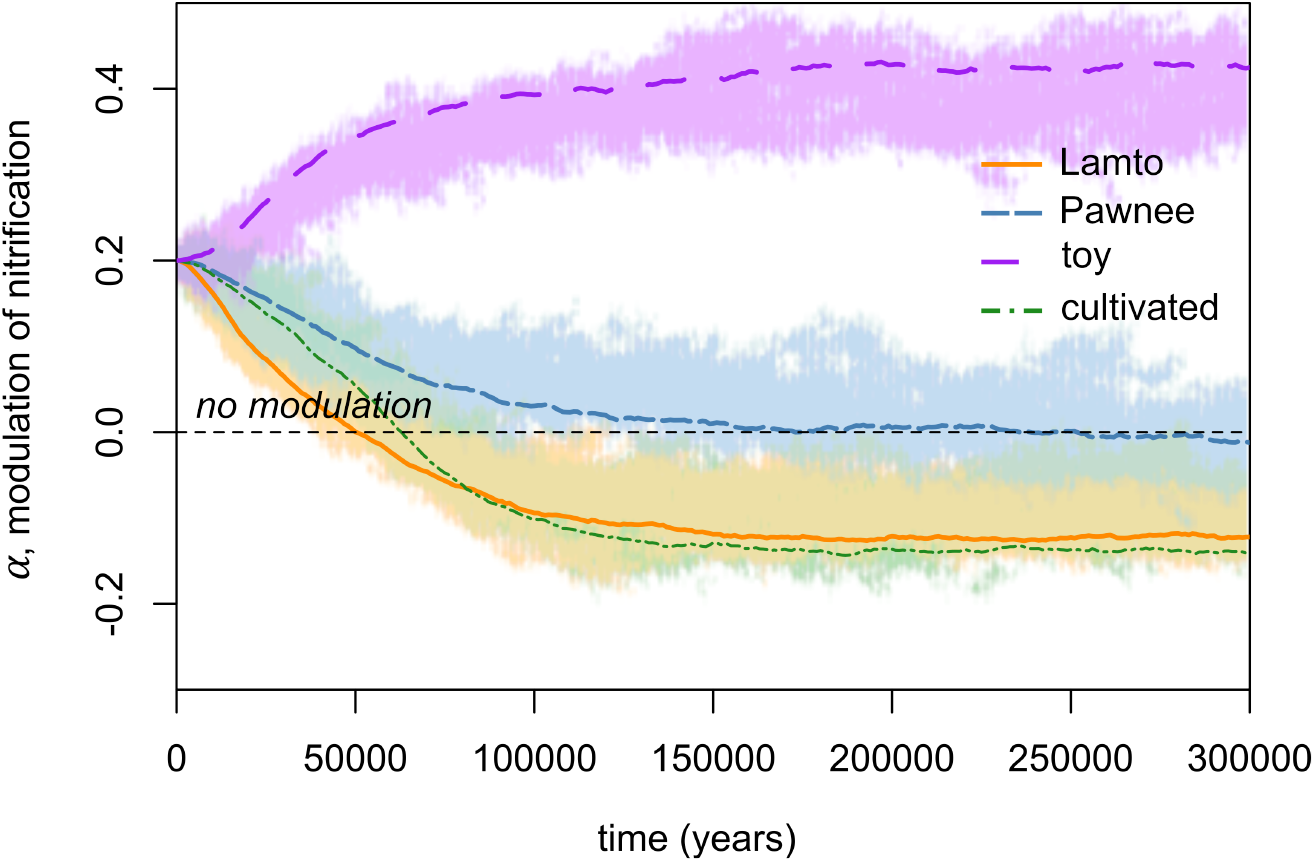
Simulations of the eco-evolutionary dynamics of plant control of nitrification with the 4 parametrizations. Pale points correspond to all strategies present on the grid for one simulation. Lines represent the mean trajectories of 10 replicates. In the Lamto savanna and cultivated system, the selected strategy is a weak inhibition. In the Pawnee prairie, the selected strategy is no control. In the toy system, the selected strategy is a strong stimulation of nitrification.

### External environmental fluxes drive the directional evolution of control of nitrification

Fig. 3a. shows that the direction of the selected control of nitrification is largely driven by loss rates and their asymmetries between ammonium and N, while the effects of N input rates are more subtle. The baseline scenario mimicking a perfectly symmetrical N cycle (equal ammonium and nitrate losses and inputs, no asymmetry in diffusion to neighboring cells), evolution drives the control of nitrification near 0. This first result refutes the hypothesis (ii-1) that inhibition should be selected for because ammonium is the first product of mineralization. In fact, in a perfectly symmetrical N cycle, short-cutting the recycling pathway does not result in higher recycling efficiency, higher plant biomass nor higher reproduction. Simulation results are however in line with our hypothesis (ii-4) that inhibition of nitrification is selected because ammonium is less mobile than nitrate. When only nitrate is mobile and diffuses to neighboring cells (*x* = 0), inhibition of nitrification is selected. Conversely, when only ammonium diffuses (*x* = 1), stimulation of nitrification is selected. Evolution of control therefore interestingly tends to allow the conservation of the most private resource. The effect of losses is as predicted by our hypothesis (ii-2): in a scenario of high nitrate losses of low ammonium losses, inhibition is selected, while low nitrate losses or high ammonium losses lead to the selection of stimulation. The effect of inputs, however, is not as we expected in hypothesis (ii-3) since the selected strategy does not change with inputs. The preference of the plant for ammonium vs. nitrate also drives the direction of the evolution of control in line with hypothesis (ii-5). Panel e shows that nitrate specialists evolve to strongly stimulate nitrification while ammonium specialists tend to evolve towards the inhibition of nitrification when leaching rates are equal.

**Figure 3:**
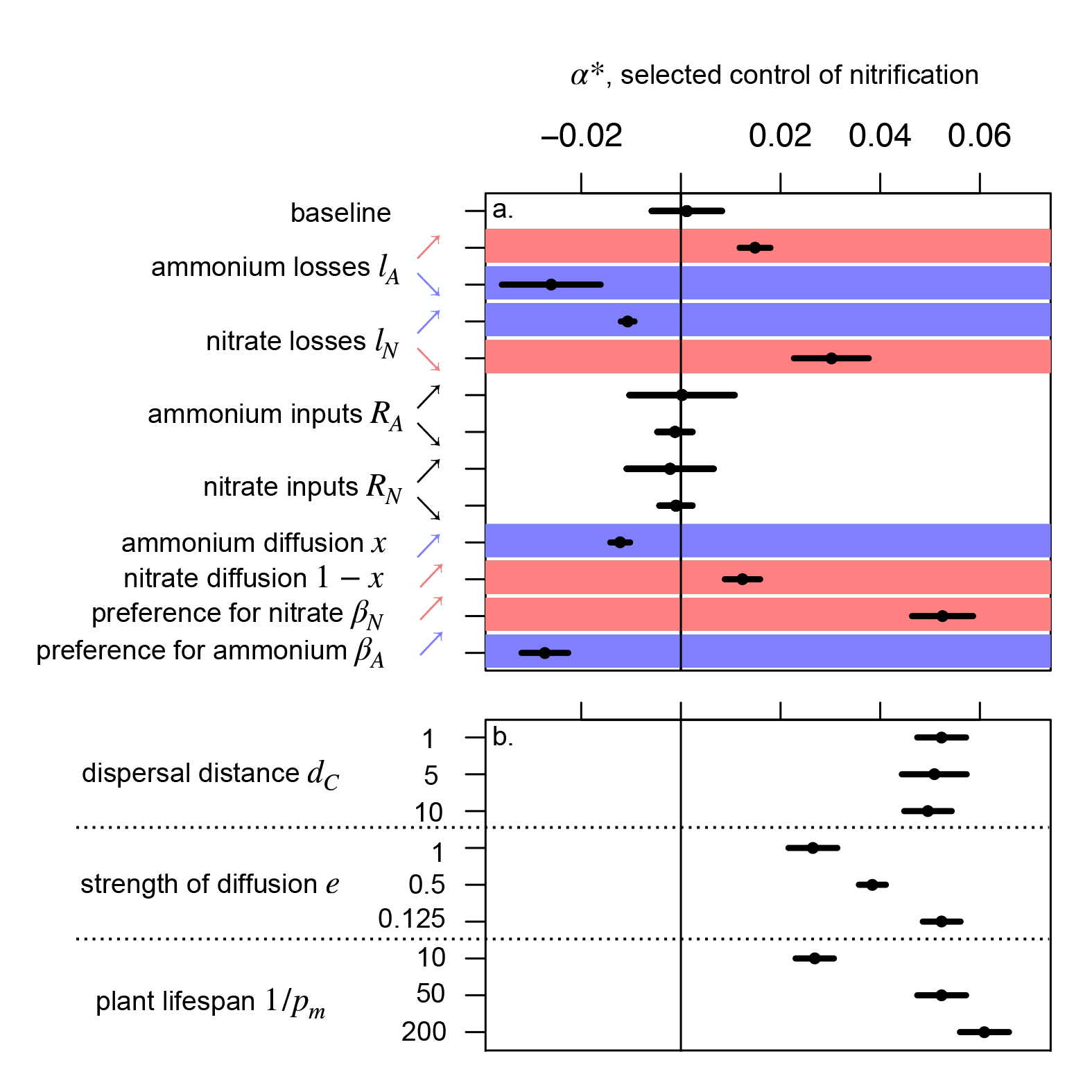
Effect of a. parameters influencing the direction of evolution and b. parameters influencing the strength of the selected control of nitrification. In a., we doubled and halved each parameter from a baseline scenario where the nutrient cycle is perfectly symmetrical. The situations leading to the selection of inhibition are highlighted in blue, the situations leading to the selection of stimulation are highlighted in red. In b., we modify three parameters that influence the strength of the selected strategy from a baseline scenario where stimulation is selected (due to strong nitrate preference, *β*_*A*_ = 0.2).

### Influence of plant life history traits and nutrient diffusion on the evolution of nitrification control

In fig. 3b., we varied two plant life history parameters (longevity and seed dispersal) and the diffusion rate of N in scenarios where stimulation has been selected. As predicted in (i-1), longer plant longevity selects for stronger control (panel g), as the niche construction effects progressively build up during the lifetime of the plant. Also coherent with the hypothesis (i-2), when the diffusion rate *e* increases, the selected control is weaker. In fact, when the diffusion rate is high, plants share their nutrients and we fall back to a ‘tragedy of the commons’ scenario where control is counter-selected (Appendix A). Finally, against the expectations formulated in (i-3), the distance of dispersal does not affect the selected control of nitrification (panel i).

### Effect of evolution on ecosystem properties

Fig. 4 illustrates that evolution may drive the plant biomass towards its maximum, or not, depending on the strength of nutrient diffusion. When nutrients are highly spatialized and do not diffuse to neighboring patches (line of squares), the selected strategy corresponds to the strategy that maximizes biomass production (squares are close to the thick white line in panel a) and minimizes total inorganic N losses (squares close to the thick white line in panel d.) This finding is in line with our previous study showing that the control of nitrification has the potential to maximize plant biomass by minimizing total N losses (Ardichvili et al., 2024b) in a mean-field model. Interestingly, the spatialization of the model does not challenge this result.

**Figure 4:**
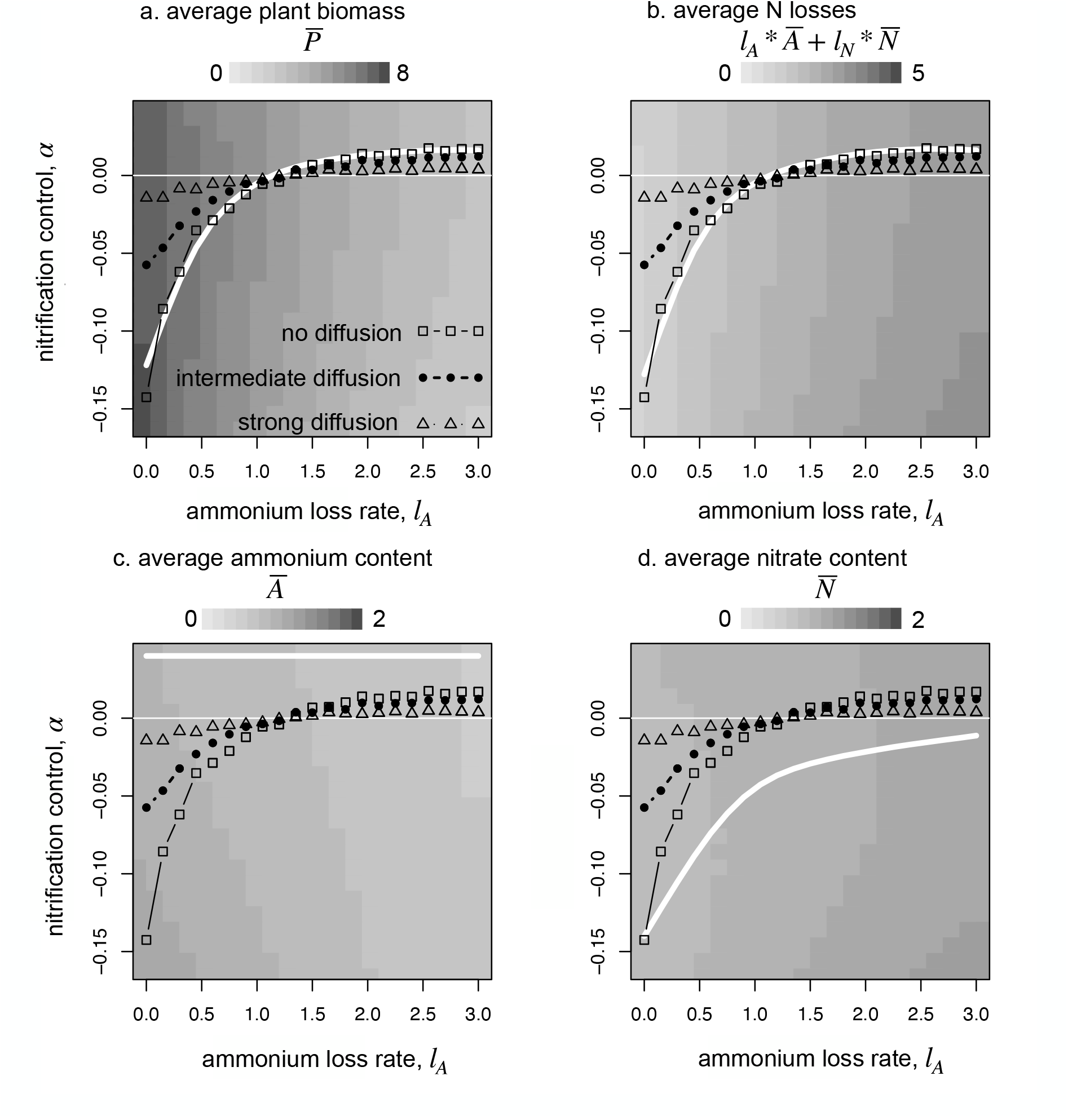
Effect of the ammonium loss rate on the selected strategy. The greyscale represents plant biomass for a fixed strategy. The thick white line illustrates the strategies maximizing plant biomass at a given ammonium loss rate in panel a, and the strategies minimizing ammonium content, nitrate content and total losses in panels b, c, d. The three black lines are the selected strategies for low (squares at e=0.125), intermediate (rounds at e=1) and strong diffusion (triangles at e=5). The diffusion rate compromises the optimization of biomass production by selection.

As the diffusion of nutrients increases (full circles and empty triangles), the selected strategy becomes weaker (ie. less nitrification control), and thus further away from the optimum biomass production. The selected strategy thus lies between the strategy that maximizes biomass production (at low diffusion rates) or the no-control strategy that maximizes the uptake rate (at high diffusion rates).

Focusing on a single nitrogen form, as in panels b and c, is not very informative in terms of understanding the link between evolution and functioning. These two panels highlight the importance of considering the losses from both nitrogen forms jointly.

## Discussion

We used a mean-field model and a spatially explicit model to study the evolution of plant capacity to control nitrification. When nutrients are shared among individuals (in the mean-field model or when the diffusion of nutrients is high in the spatially explicit model), control of nitrification is counter-selected. In the spatially explicit model, each plant individual has greater control of its own nitrogen cycle at the rhizosphere level, as reported in empirical studies (Subbarao et al., 2006). When the environment of the plant is sufficiently private (i.e. when the diffusion of nutrients to neighboring cells is sufficiently low), activation or inhibition can be selected depending on plant preference for either form, which inorganic form is the shared resource, and relative losses of ammonium and nitrate. When ammonium losses are lower than nitrate losses, plants evolve to inhibit nitrification; when nitrate losses are higher than ammonium losses, plants evolve to stimulate nitrification. Our results suggest that global studies aiming to characterize the control strategy of plants (such as the GainGRASS project) should also estimate these key environmental parameters (loss rates of ammonium vs. nitrate, deposition rate, and preference) to interpret the geographic distribution of nitrification-inhibiting or -stimulating plants.

Our results show that nitrification control is not selected in the absence of spatial structure. This is a classical tragedy of the commons scenario (Hardin, 1968). Controlling nitrification is costly for the plant individual, while the benefits are diluted in the whole population as all nutrients are shared. In a population of non-controlling plants, a controlling mutant bears the cost on the uptake rate but since it is rare, its effect on the nitrification rate is weak; its growth rate is lower than the resident and it cannot invade the population. In a population of controlling plants, a non-controlling (or less controlling) mutant benefits from the controlling activity of conspecifics without paying any cost. Its growth rate is higher and it can invade the population.

The spatial version of the model solves this tragedy of the commons. Evolution towards inhibition or stimulation occurs, in line with hypotheses (i-1 and i-2), when plant longevity is sufficient (low *p*_*m*_) and when the diffusion of N forms is limited (low *e*). When the diffusion of nutrients is limited, plants experience a private environment and the direct benefits of control become larger for the plants. Such a result echoes those of a more general model by Kylafis and Loreau (2008) showing that a niche construction trait bringing both direct benefits to the constructing individual and benefits shared by the whole population, could only evolve when direct benefits were sufficient. It also echoes empirical observations. For example, in the Lamto savanna, BNI grasses such as *Andropogon schirensis, Andropogon canaliculatus* or *Hyparrhenia diplandra* grow in tussocks, with large areas of bare soil in between (Lata et al., 1999; Srikanthasamy et al., 2018). The tussock life form, possibly an adaptation to savanna fires (Archibald et al., 2019), may also have promoted the evolution of the control of nitrification because individuals have a local private environment. In turn the local constructed nitrogen conditions can create suitable local conditions that favor the tussock development and maintenance. We predict that perennial grasses that do not grow in tussocks, such as *Cynodon dactylon*, may not have evolved to control nitrification.

Having a long lifespan is a way to experience a private environment through time, which subsequently favors the evolution of nitrification control. At the time of the establishment, when plant biomass is low, plants strongly experience the cost of control, decreasing their growth rate. It takes time for plant biomass to build up sufficiently for the root exudates to impact the N cycle and for the local availability of mineral nitrogen to build up. When the lifespan of plants is short, controlling individuals only experience the cost of the control, their growth rate is diminished and such plants are counter-selected. When the lifespan of plants is long, niche construction effects progressively build up and plants experience the benefits associated to nitrification control. This result is in line with the fact that inhibiting grasses in the Lamto savanna are perennialand live on average 80 years (Koffi et al., 2019). The model predicts that it is unlikely to find nitrification controlling annual or biannual plants since the control should be counter-selected if costly. This result is consistent with Grime (1977)’s idea that long-lived plants are stress-tolerating species that have evolved strategies enabling the conservation of resources, as opposed to short-lived, competitive, species which have evolved strategies of high resource acquisition.

Contrary to expectations, the dispersal trait of the plant did not affect the mean selected control strategy, which was the same for short and long dispersal ranges. The fact that low dispersal ranges does not promote the evolution of control is unexpected given the role of aggregation for the evolution of altruistic traits (Ferrière and Le Gaillard, 2001; Lion and van Baalen, 2008; Ross-Gillespie et al., 2007). In two spatially explicit previous models of nutrient cycling, a low dispersal distance was key to allow the evolution of two traits susceptible to the tragedy of the commons (evolution of low uptake rates Barot et al., 2016, and evolution of the ability to increase mineralization rates Barot et al., 2014). The evolution of control, even when strategies are randomly distributed (when the dispersal distance is large), questions the nature of the interaction between nitrification-controlling individuals. Controlling individuals consume a large fraction of the N form that is kept locally due to the control of nitrification (Appendix C). Since empty cells are richer than occupied cells whether the occupant is a controlling or non-controlling individual, the inhibition (resp. stimulation) of nitrification by the neighbor does not result in a subsidy flow of ammonium (resp. nitrate) to the empty cell at the time of establishment. The control of nitrification merely mitigates the competition between neighbors but does not result in facilitation between neighbors. We previously showed that the conditions resulting in facilitation between organisms consuming resources, even when niche construction modifies these resource flows, are limited (Ardichvili et al., 2024a).

When the evolution of nitrification control was possible, the effects of asymmetry in diffusion and inorganic losses are consistent with hypotheses (ii-2,4). In a system where losses are increased, plants evolve to control nitrification in the direction that decreases the residence time of N in the compartment with the highest losses. This result is very much in line with the hypotheses proposed by many empiricists as to why biological inhibition should have been selected in many ecosystems where nitrate losses as leachates are much more important than ammonium losses (Lata et al., 2022; Rice and Pancholy, 1972; Subbarao et al., 2006). The model predicts that in ecosystems where ammonium is more mobile (which could be the case of ecosystems with an acidic litter in which ammonium is readily lost by volatilization, Craig and Wollum, 1982), stimulation of nitrification should be selected. A common garden experiment showed that some coniferous trees (typically producing an acidic litter) stimulate nitrification (Andrianarisoa et al., 2010).

In these simulations, we considered that the preference of plants was fixed (to the generalist strategy in the baseline scenarios, *β*_*A*_ = *β*_*N*_ = 0.5) and predict that the control of nitrification should evolve towards the increase in the preferred form of mineral nitrogen. A previous simulation study indeed suggested that a strong preference for ammonium should be selected for in a savanna ecosystem (Boudsocq et al., 2012). It would be interesting to test whether the coevolution of nitrification (niche construction trait or effect trait, *sensu* Lavorel and Garnier, 2002) and preference (niche trait or response trait) would alter the evolutionary outcome.

In our spatially explicit model, the selected control of nitrification tends towards the level of control that maximizes biomass (and minimizes total N losses) when the diffusion is weak. None of the other parameters affecting the strength of the selected control (mortality rate, strength of the trade-off) affected this optimization. Our study provides an example where, depending on the level of privatization of the resources, evolution optimizes the abundance (at low diffusion rates) or optimizes the growth rate (at high diffusion rates). In natural ecosystems, such horizontal flows of nutrients can be driven by plant ability to forage outside of their rhizosphere by forming mycorrhizal associations. The level of privatization of the two nitrogen forms depends on the diffusion capacity of the two ions, and also to some extent on plant root traits. In general, situations of biomass optimization or growth rate optimization do not allow the evolutionary divergence of strategies (Metz et al., 2008; Rohr and Loeuille, 2023). Here, in all simulations, the control converged to a single mean trait, although large variation around the mean were possible: there is no sympatric diversification of the control strategy This result apparently contradicts field observations (e.g. Lamto savanna), where inhibiting grasses coexist with stimulating trees, both strategies forming patches (Srikanthasamy et al., 2018) and where grasses with different control strategies coexist (Yé et al., 2015). Our results suggest that environmental parameters exert a strong influence on the selected control. We modeled a homogeneous biome in which loss rates and inputs are constant. Trees and grasses possibly experience different loss rates, trees being able to forage deeper in the soil (Konaré et al., 2019). Introducing variability in loss and input rates (for example at the landscape scale by modeling higher nitrate losses at the top of a slope) could promote the evolutionary divergence of strategies. The diversification of the control strategy could evolve in allopatry but not in sympatry.

## Conclusion

To conclude, we showed that control of nitrification can evolve whenever plants are sufficiently long-lived and when their nutrient stocks are sufficiently private. There are potentially many species (any displaying those characteristics) that could have evolved the ability to control nitrification, which should incite empirical measures of such capacities in wild species. The ability to control nitrification, as other niche constructing traits, could be integrated to studies aiming to classify plant strategies or place them on a Root/Leaf economic spectrum (Weigelt et al., 2021). Our model does not intend to make quantitative predictions. However, our qualitative results suggest that understanding nitrogen cycling and predicting N_2_O emissions through denitrification (that are directly linked to nitrification and that have a strong greenhouse gas effect) at the local and global scales requires considering the evolution of the control of nitrification by plants. This would be key to take into account feedbacks between vegetation and climate. Finally, inhibition of nitrification is currently thought of as a way to improve the N efficiency of cultivated systems (Subbarao and Searchinger, 2021), minimize N leaching and nitrous oxide emissions. The artificial selection of BNI varieties requires evolutionary considerations. For example, our model suggests strategies to find plant species that are inhibiting nitrification due to the constraints of their environment. Our results also suggest that it would be critical to asses the costs linked to the production of nitrification-inhibiting exudates to determine whether the selection of nitrification-inhibiting crops is likely or not to increase biomass production or the crop yield. Our model suggests that the selection of BNI capacity should be possible in perennial cereals, which are currently selected and being considered as more sustainable crops (Cox et al., 2006).

## Acknowledgments

This work was granted access to the HPC resources of the SACADO MeSU platform at Sorbonne Université. We thank all members of the GainGRASS project (Global Assessment of Nitrification Inhibition by tropical Grasses Project, https://anr.fr/Project-ANR-19CE02-0009). JCL and SB were supported the GainGRASS project, AA was funded by a doctoral grant from the French research ministry.

## Appendix A: Adaptive dynamics of the control of nitrification in a mean-field model

### Obtaining the invasion fitness

From the ecological model (Eq. 1), we derive the dynamics of a dominant population *P* interacting with a mutant population *P*_*m*_ with trait *α*_*m*_.

The main assumption of adaptive dynamics is the separation of evolutionary and ecological timescales. The introduced mutant is assumed to be rare (*P*_*m*_ ≈ 0) while the resident has reached equilibrium (*P*^*\**^). Using these two assumptions, we write the invasion fitness of a mutant, that is, the *per capita* growth rate of a rare mutant with trait *α*_*m*_ in the stationary conditions set by the resident population of trait value *α* :

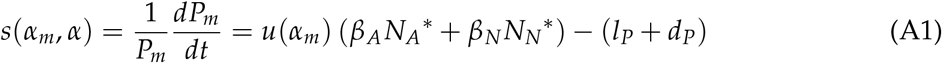

with

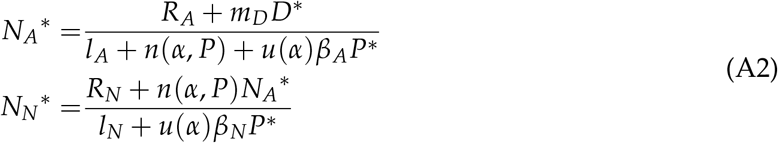

We use the invasion fitness (Eq. A1) to find the long-term evolutionary outcome (evolutionary singularity) and its second derivatives to compute the convergence and invasibility criteria (Geritz et al., 1998).

From Eq. 1, when the resident is at equilibrium (*dP*^*\**^ /*dt* = 0), we write:

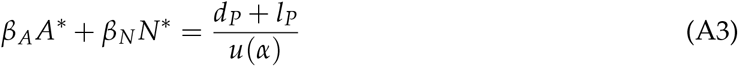

Replacing Eq. A3 into the invasion fitness equation Eq. A1, we get Eq. 6.

#### Convergence and invasibility

From equation 6 we can write the local fitness gradient:

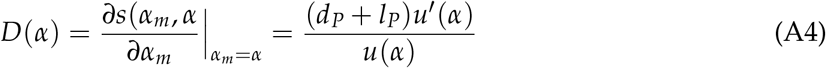

Evolution follows the fitness gradient and stops the fitness gradient becomes null. In this case, since the cost function *u*(*α*) is maximal when plants do not control nitrification; the evolutionary singularity (ES) is *α*^*\**^ = 0.

The sign of the second derivative at the singularity determines whether the singularity is invasible or not. Here:

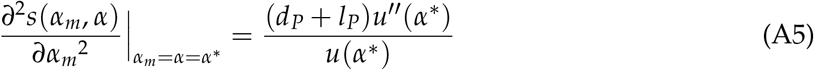

Since 0 is a maximum for *u*(*α*), *u*^*′′*^(*α*^*\**^) is negative and the ES is ESS-stable (non-invasible). Whether the ES is convergent depends on the sign of:

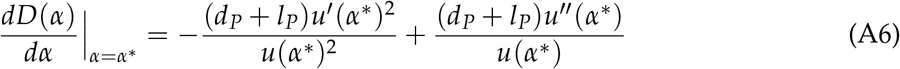

The first term is null and the second term is negative. The ES is convergent.

The ES being convergent and non-invasible, *α*^*\**^ = 0 is a Continuously Stable Strategy.

## Appendix B: Effect of the selected strategy on regional ammonium and nitrate content, and losses

In this appendix, we show the influence of evolution on ecosystem properties other than plant biomass: inorganic N content and total inorganic N losses. The figure illustrates that for every parameter we varied, the selected strategy maximizes biomass, while the effect on ammonium and nitrate content is more subtle. However, calculating the total inorganic N losses as *A*^*\**^ *× l*_*A*_ + *N*^*\**^ *× l*_*N*_, we see that the selected strategy minimizes total N losses. This finding is in line with our previous study showing that the control of nitrification has the potential to maximize plant biomass by minimizing total N losses (Ardichvili et al., 2024b) in a mean-field model. Interestingly, the spatialization of the model required to observe the evolution of nitrification control does not challenge this result.

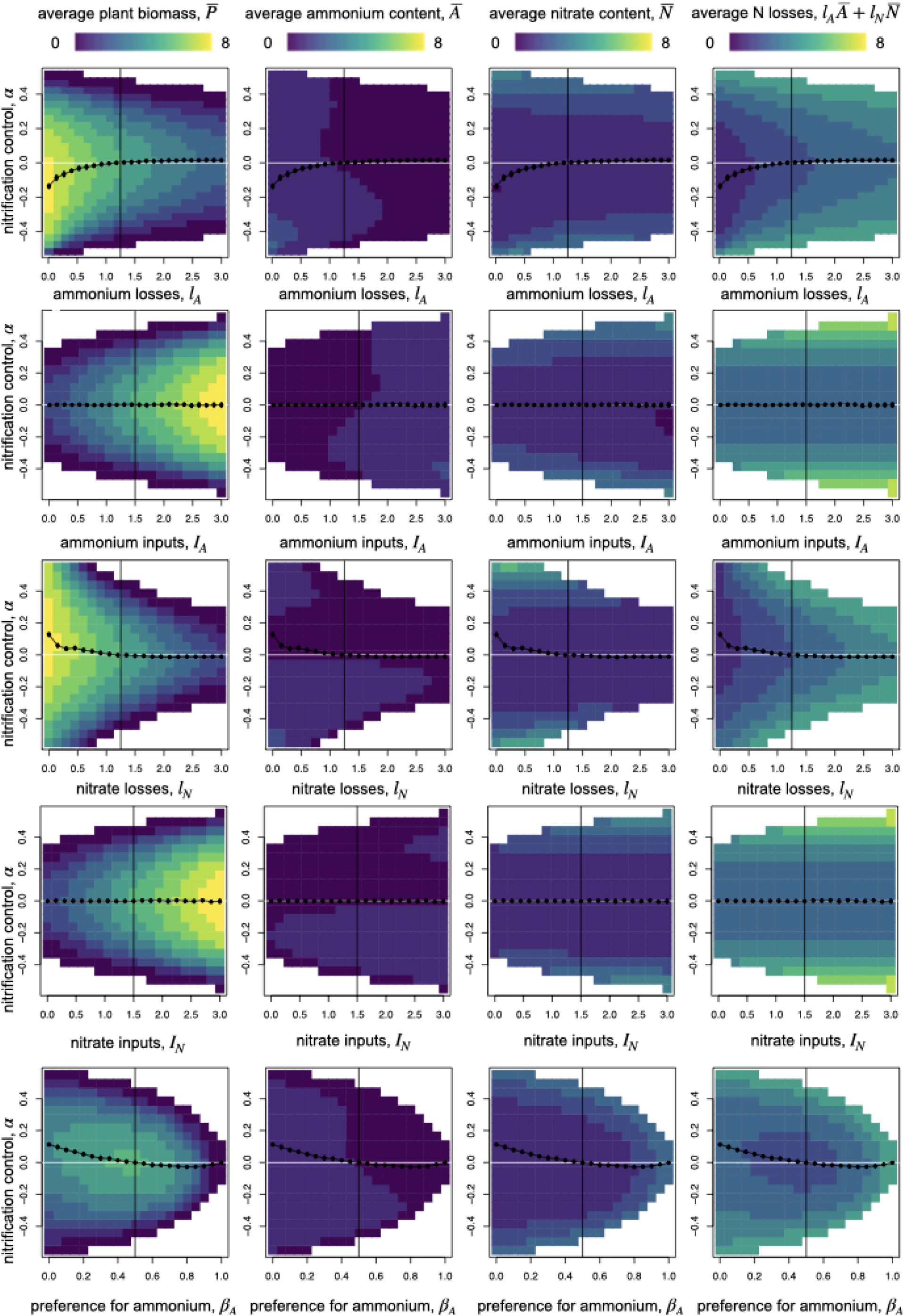

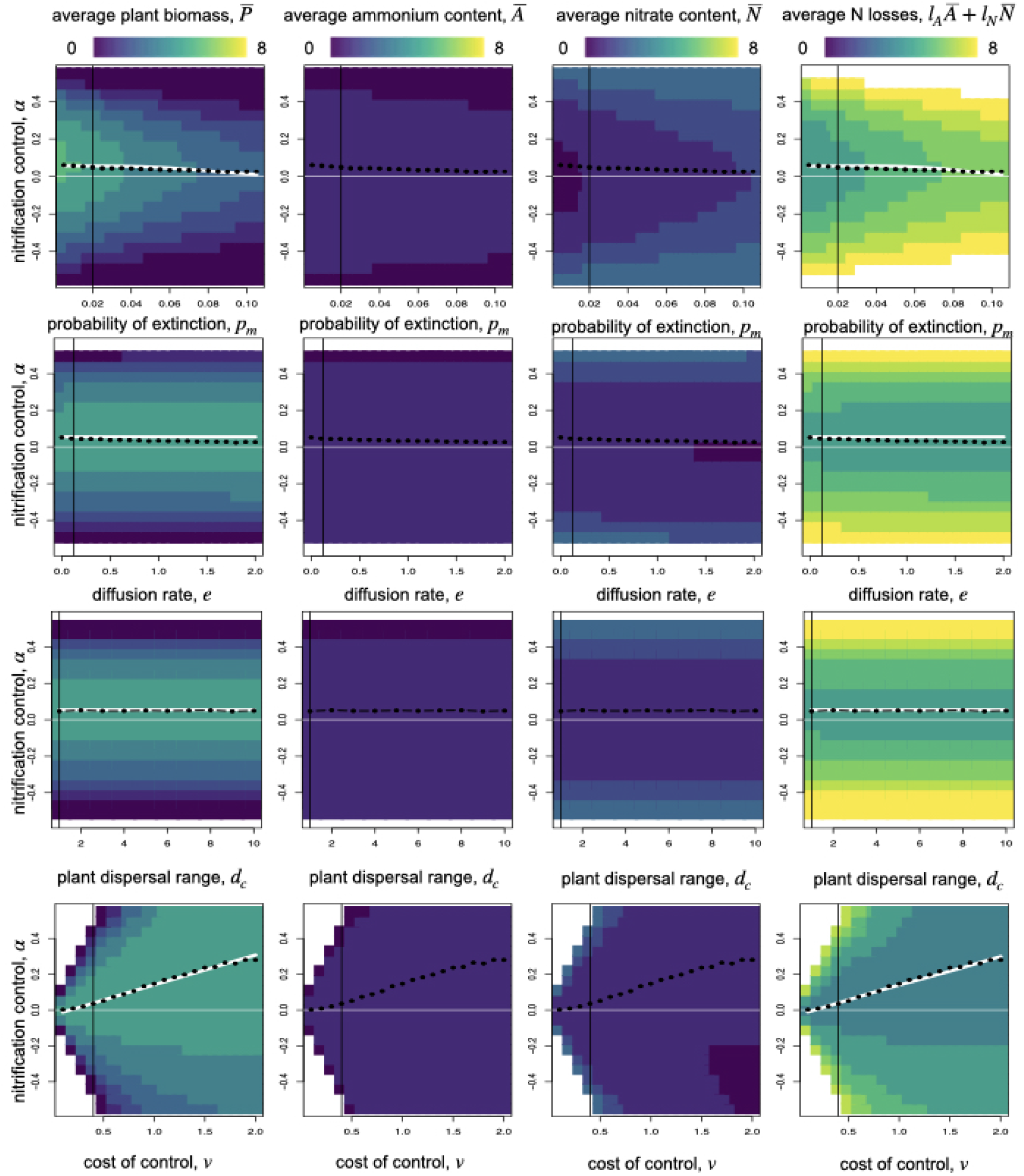

## Appendix C: Control of nitrification mitigates competition between neighbors

We predicted that control traits, i.e. niche creating phenotypes, should be aggregated for control to evolve. Such aggregation is achieved when the dispersal of phenotypes is low, while a high dispersal breaks down the aggregation of phenotypes. Contrary to our expectations, we find that control of nitrification can evolve even when the dispersal distance is large. This finding questions the extent to which controlling individuals should be considered as ‘cooperators’ or as ‘competitors’.

Fig. C1 illustrates that plant individuals first destroy their niche by consuming the locally available nutrient. In panels A and B, we see two clusters of points, with a few scattered in the middle of them. The cluster in the top left of the figure corresponds to empty cells or cells that have just been colonized, in which plant N content is low but inorganic N content (be it ammonium or nitrate) is high. The cluster in the bottom right corresponds to occupied cells that have reached their equilibrium biomass (high N content in plants) and consumed the available nutrient (low ammonium and nitrate content). Within this cluster of points, cells occupied by strongly inhibiting individuals are richer in ammonium than those occupied by less inhibiting individuals (see the succession of layers of black (strong inhibitors), purple (inhibitors) and orange (mild inhibitors) points within the cluster in A.). The effect on the richness of the cell in terms of nitrate is the same regardless of the strategy (points in the bottom right does not show any layering in B.). The Fig. C1 illustrates two things: occupied cells are always poorer in terms of ammonium than empty cells, implying that the interaction between two neighboring plants is competition. Second, strongly inhibiting plants reduce ammonium content in their cell less than less controlling counterparts. Therefore, the competition between two strongly inhibiting plants should be less intense than between two less-inhibiting plants.

**Figure C1:**
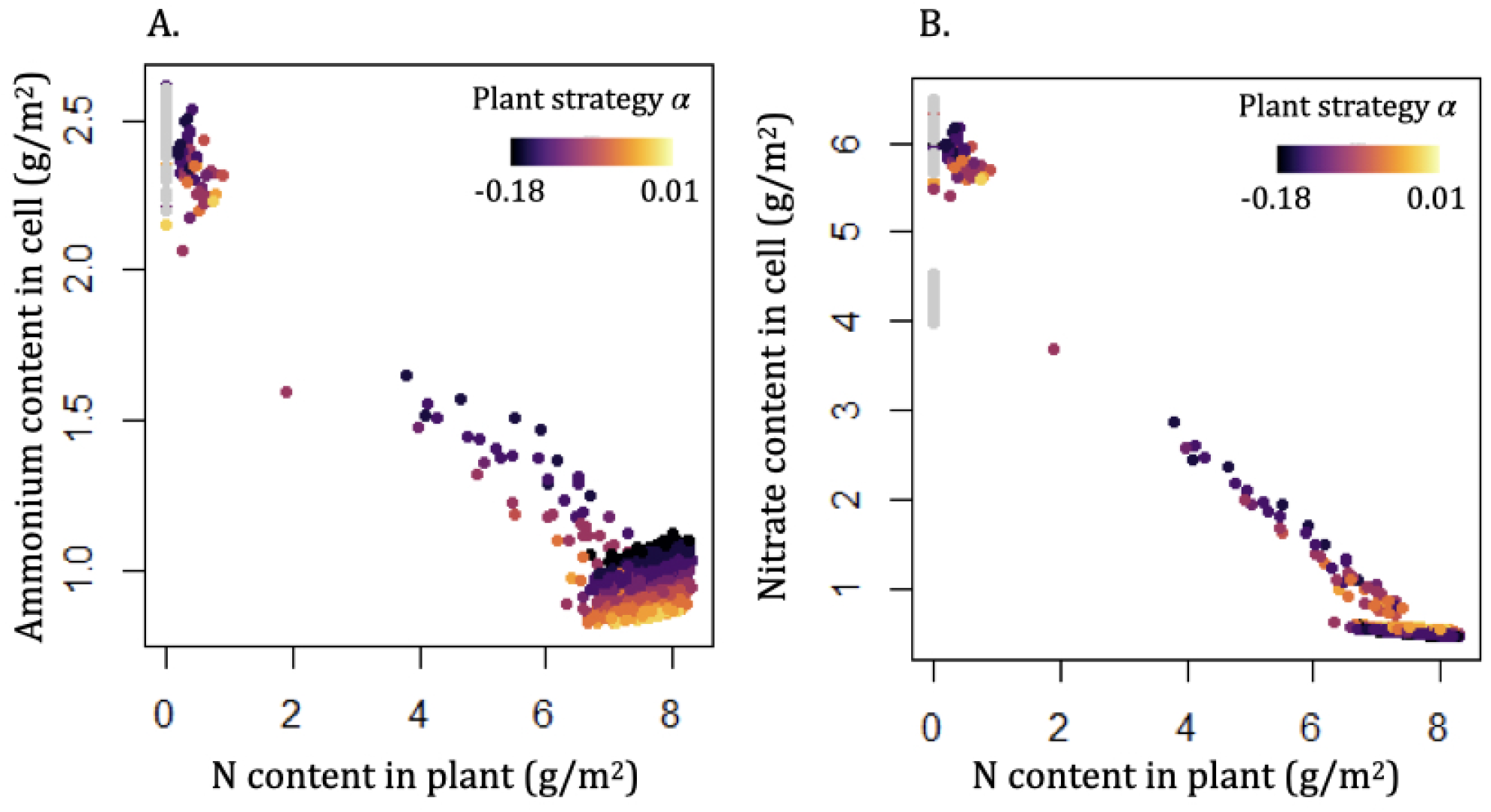
Relationship between plants N content, and ammonium (A) or nitrate (B) content in the cell. Each dot represents a cell at the end of a simulation with low dispersal, low diffusion, and weak ammonium losses (*d*_*C*_ = 1, *e* = 0.1, *l*_*A*_ = 0.0625, *l*_*N*_ = 1.25, *R*_*A*_ = 1.5, *R*_*N*_ = 1.5). The color of the dots depends on the plant control trait in the cell. The figure for long-range dispersal (*d*_*C*_ = 10) is not shown, but displays the same pattern.

